# Targeting both GD2 and B7-H3 using bispecific antibody improves tumor selectivity for GD2-positive tumors

**DOI:** 10.1101/2024.05.23.595624

**Authors:** Zachary T. Rosenkrans, Amy K. Erbe, Nathan B. Clemons, Arika S. Feils, Yadira Medina-Guevara, Justin J. Jeffery, Todd E. Barnhart, Jonathan W. Engle, Paul M. Sondel, Reinier Hernandez

## Abstract

**Objectives:** Disialoganglioside 2 (GD2), overexpressed by cancers such as melanoma and neuroblastoma, is a tumor antigen for targeted therapy. The delivery of conventional IgG antibody technologies targeting GD2 is limited clinically by its co-expression on nerves that contributes to toxicity presenting as severe neuropathic pain. To improve the tumor selectivity of current GD2-targeting approaches, a next-generation bispecific antibody targeting GD2 and B7-H3 (CD276) was generated.

**Methods:** Differential expression of human B7-H3 (hB7-H3) was transduced into GD2^+^ B78 murine melanoma cells and confirmed by flow cytometry. We assessed the avidity and selectivity of our GD2-B7-H3 targeting bispecific antibodies (INV34-6, INV33-2, and INV36-6) towards GD2^+^/hB7-H3^-^ B78 cells relative to GD2^+^/hB7-H3^+^ B78 cells using flow cytometry and competition binding assays, comparing results an anti-GD2 antibody (dinutuximab, DINU). The bispecific antibodies, DINU, and a non-targeted bispecific control (bsAb CTRL) were conjugated with deferoxamine for radiolabeling with Zr-89 (t_1/2_ = 78.4 h). Using positron emission tomography (PET) studies, we evaluated the in vivo avidity and selectivity of the GD2-B7-H3 targeting bispecific compared to bsAb CTRL and DINU using GD2^+^/hB7-H3^+^ and GD2^+^/hB7-H3^-^ B78 tumor models.

**Results:** Flow cytometry and competition binding assays showed that INV34-6 bound with high avidity to GD2^+^/hB7-H3^+^ B78 cells with high avidity but not GD2^+^/hB7-H3^+^ B78 cells. In comparison, no selectivity between cell types was observed for DINU. PET in mice bearing the GD2^+^/hB7-H3^-^ and GD2^+^/hB7-H3^+^ B78 murine tumor showed similar biodistribution in normal tissues for [^89^Zr]Zr-Df-INV34-6, [^89^Zr]Zr-Df-bsAb CTRL, and [^89^Zr]Zr-Df-DINU. Importantly, [^89^Zr]Zr-Df-INV34-6 tumor uptake was selective to GD2^+^/hB7-H3^+^ B78 over GD2^+^/hB7-H3^-^ B78 tumors, and substantially higher to GD2^+^/hB7-H3^+^ B78 than the non-targeted [^89^Zr]Zr-Df-bsAb CTRL control. [^89^Zr]Zr-Df-DINU displayed similar uptake in both GD2^+^ tumor models, with uptake comparable to [^89^Zr]Zr-Df-INV34-6 in the GD2^+^/hB7-H3^+^ B78 model.

**Conclusion:** The GD2-B7-H3 targeting bispecific antibodies successfully improved selectivity to cells expressing both antigens. This approach should address the severe toxicities associated with GD2-targeting therapies by reducing off-tumor GD2 binding in nerves. Continued improvements in bispecific antibody technologies will continue to transform the therapeutic biologics landscape.

**Graphical Abstract:** 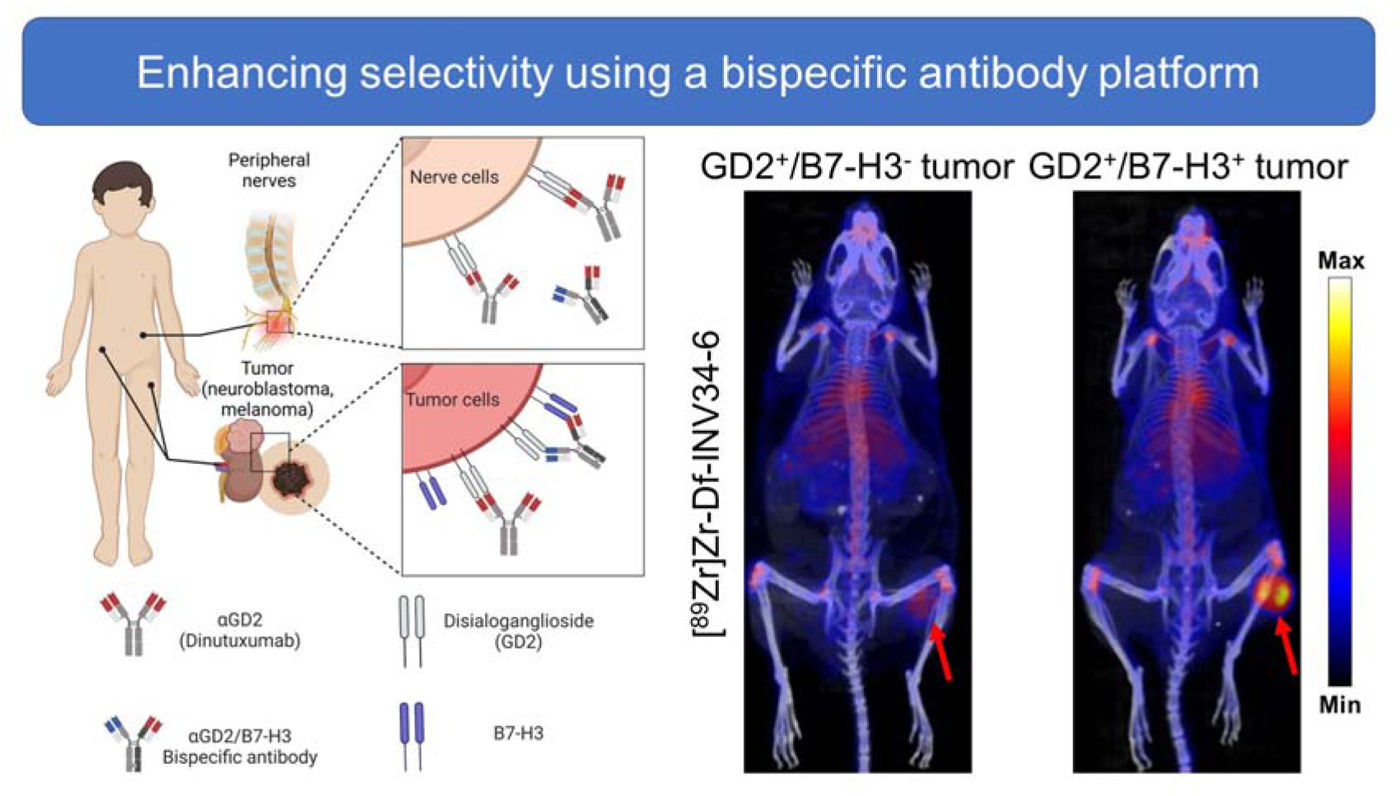

## Introduction

Antibodies have been developed as immunotherapies of cancer by targeting disialoganglioside 2 (GD2), which is highly expressed on the surface of solid tumors such as neuroblastomas or melanomas.[1–3] Patients treated with anti-GD2 immunotherapy (dinutuximab, DINU) experience neuropathic pain arising from GD2 expression on peripheral nerve fibers.[4] The pain caused by anti-GD2 immunotherapy is ubiquitous across patients, can be debilitating, and narrows the therapeutic index.[5, 6] B7-H3 (CD276), an immune checkpoint antigen with limited expression on normal tissues such as liver and some immune cells, is highly expressed on many tumors, and is not expressed on nerves.[7–9] As such, immunotherapies targeting B7-H3 have been well tolerated with some anti-tumor activity in clinical trials.[10, 11] Thus, improving cancer cell selectivity by requiring co-expression of B7-H3 together with GD2 may potentially mitigate the agonizing pain induced by GD2-directed monoclonal antibodies.

Therapeutic immunoglobulin G (IgG) antibodies are emerging as the predominant treatment modality due to their pharmacokinetic properties, high specificity, and effector-driving function.[12] Advances in protein engineering techniques have facilitated the development of bispecific antibodies (bsAbs) that target two antigens or epitopes. The introduction of bsAbs has transformed the biological drug landscape, presenting a novel class of agents with therapeutic or diagnostic potential beyond the capabilities of monospecific IgG counterparts. The innovation of bsAbs permits novel strategies through additional control of avidity, specificity, biodistribution, and therapeutic mechanism of action. The emergence of bsAbs has spurred extensive clinical investigation, leading to successful regulatory approval of several.[13, 14]

We aim to improve the precision of antibody targeting toward GD2-positive tumors using a bsAb design that targets both GD2 and B7-H3 antigens. Our next generation approach leverages a requirement of GD2 and B7-H3 co-expression to enhance cancer cell selectivity, minimizing binding and side effects associated with anti-GD2 mAb binding to GD2 expressed by nerves. Through positron emission tomography (PET), we evaluated the avidity and selectivity of GD2/B7-H3 bsAbs *in vivo*. By requiring the simultaneous presence of the GD2 and B7-H3 on the same cell, we anticipate an improved antibody agent that selectively targets tumor that co-express GD2 and B7-H3 but not to nerves that express GD2 but not B7-H3.

## Materials and methods

### Antibody conjugation for PET

Antibodies used in this study were kindly supplied by Invenra Inc (Madison, WI). p-SCN-Bn-Deferoxamine (Df) from purchased from Macrocyclics (Plano, Tx) was conjugated to antibodies for PET imaging. Using isothiocyanate chemistry, Df was dissolved in anhydrous DMSO and conjugated antibodies in PBS pH adjusted to 8.5-9 at a molar ratio of 5:1 (Df:antibody). Using end over end rotation, the reactions were conducted for 2-4 h at room temperature. Df conjugated antibodies were purified using a Cytiva (Marlborough, MA) PD-10 size exclusion chromatography column

### Antibody radiolabeling

The UW-Madison Cyclotron Group produced [^89^Zr]Zr-oxalate using the ^nat^Y(p,n)^89^Zr reaction on a PETtrace cyclotron (GE Healthcare [Madison, WI]). Zr-89 was used to radiolabel Df conjugated antibodies. For radiolabeling, Df conjugated antibodies were added to Zr-89 (100 ug per 37 MBq [1 mCi]) in 1 M HEPES buffer. Typical reactions utilized 1-2 mCi (37-74 MBq) of Zr-89. Radiolabeling reactions were performed at 37°C for 1 h and purified using a PD-10 size exclusion chromatography column (Cytiva). Instant thin layer chromatography (iTLC) was used to determine the radiolabeling efficiency.

### Animal studies

All animal studies were conducted on a protocol approved by the Institutional Animal Care and Use Committee at the UW-Madison. Female or male C57BL/6 were purchased from Taconic and housed at the UW-Madison.

### B78 tumor model

To establish B78 tumor allografts, mice were subcutaneously injected with 2x10^6^ B78 cells engineered with differential GD2 and B7-H3 expression (GD2^+^/hB7-H3^-^ or GD2^+^/hB7-H3^+^). GD2^+^/hB7-H3^-^ B78 cells were established by CRISPR Cas9 knockout of endogenous murine B7-H3 in the GD2^+^ B78 parental cell line. GD2^+^/hB7-H3^+^ B78 cells were established by lentivirus transduction of hB7-H3. Imaging studies utilized GD2^+^/hB7-H3^+^ B78 and or GD2^+^/hB7-H3^-^ tumors implanted on the lower right flank of mice.

### PET/CT imaging studies

Following intravenous injection of approximately 3.7 MBq – 7.4 MBq (100 µCi – 200 µCi) of radiotracer, PET or PET/CT were acquired using an Inveon µPET/CT. CT images were captured using the following parameters: 80 kV, 900 µA, resolution of 105 µm. PET/CT images were acquired a3 h, 24 h, 48 h, or 72 h post injection and reconstructed using an OSEM3D/MAP algorithm. Volume of interest (VOI) quantification of the collected images was performed in the Inveon Research Workstation. VOI quantified tissues included the blood, liver, spleen, tumor, kidney, muscle, and bone, reported in percent injected activity per cubic centimeter of tissue (%IA/cc). Ex vivo quantification of the injected mice was performed following the final imaging time point by harvesting tissues. A Wizard 2 (Perkin Elmer [Waltham, MA]) or Hidex AMG (Hidex [Turku, Finland]) gamma counter was used to quantify radioactivity. Quantification of the ex vivo biodistribution data was reported in percent injected activity per gram of tissue (%IA/g).

### Statistical Analysis

Statistical analysis was performed by two-tailed unpaired Student’s *t* tests or one-way ANOVA. NS, non-significant \**P*__<__0.05, \*\**P*__<__0.01, \*\*\**P*__<__0.001.

## Results

### Designing a bispecific antibody platform targeting GD2 and B7-H3 using a bispecific antibody approach

Bispecific antibodies targeting GD2 and B7-H3 were prepared (**Fig. 1A**) in an IgG like format using the “knob-in-hole” mutations in the heavy chain constant region three [15, 16] with one arm containing a B7-H3 binding domain and one arm containing a GD2 binding domain. The selectivity of the bispecific antibody was designed by targeting B7-H3 with high-medium affinity and GD2 with low affinity in their respective Fab variable regions. We anticipated that the combination of these affinities should generate an bispecific antibodies that selectively target GD2^+^/hB7-H3^+^ cells with high avidity (**Fig. 1B**).

**Figure 1.**
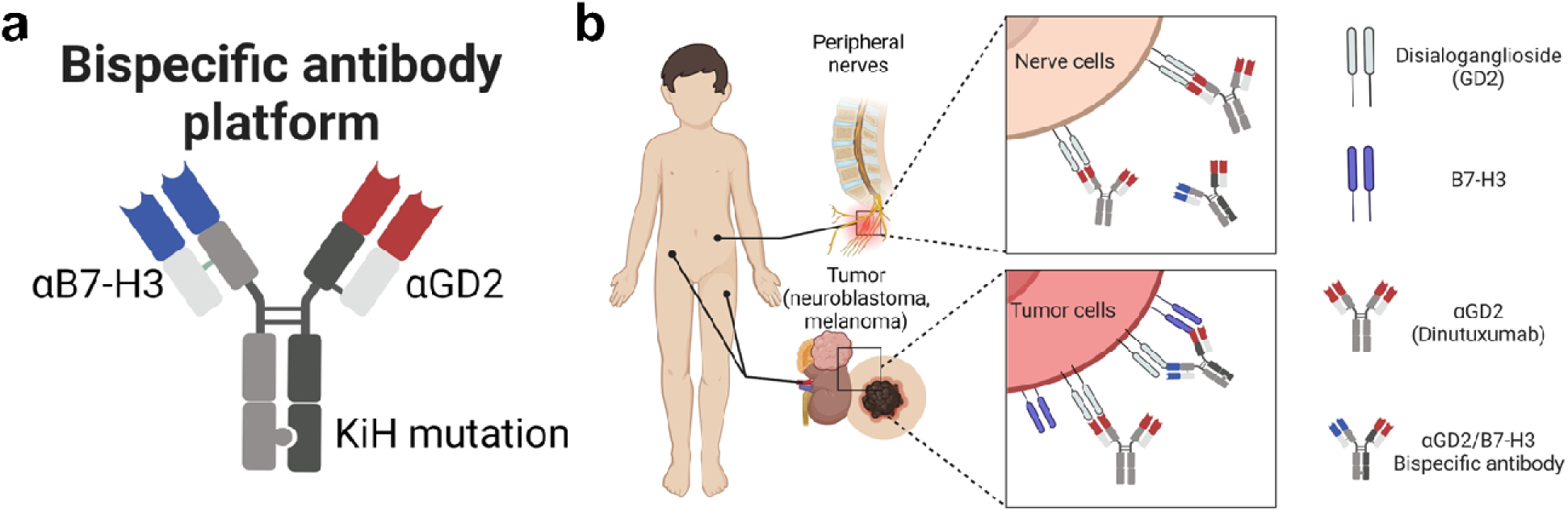
A bispecific antibody platform that enables next generation targeting of tumors. (**a**) Bispecific antibody construct with one arm containing a B7-H3 binding domain (αB7-H3), one arm containing a GD2 binding domain (αB7-H3), and an Fc containing the knob-in-hole (KiH) mutation (**b**) Bispecific antibody selectively targets GD2^+^/hB7-H3^+^ tumor cells with high bivalent avidity (upper panel) compared to low affinity monovalent binding to GD2^+^/hB7-H3^-^ nerve cells (bottom panel).

### Screening in vitro avidity and selectivity for the bispecific GD2/hB7-H3 targeting antibody

The avidity and selectivity of the GD2/hB7-H3 targeting bispecific antibodies were screened using flow cytometry (**Fig. 2A**). Using GD2^+^ B78 cells that express the human variant of B7-H3 (hB7-H3), bispecific antibody INV34-6 bound with high avidity when both antigens were present on the cells (EC_50_ = 3.1 nM). In comparison, negligible binding of INV34-6 to B78 cells expressing GD2 only (simulating nerve) was observed (EC > 100 nM). As shown in **Fig. 2B**, we then evaluated DINU binding and found that DINU bound with high and comparable affinity to GD2^+^/hB7-H3^-^ (EC_50_ = 13.5 nM) or GD2^+^/hB7-H3^+^ (EC_50_ = 29.9 nM) B78 cells.

**Figure 2.**
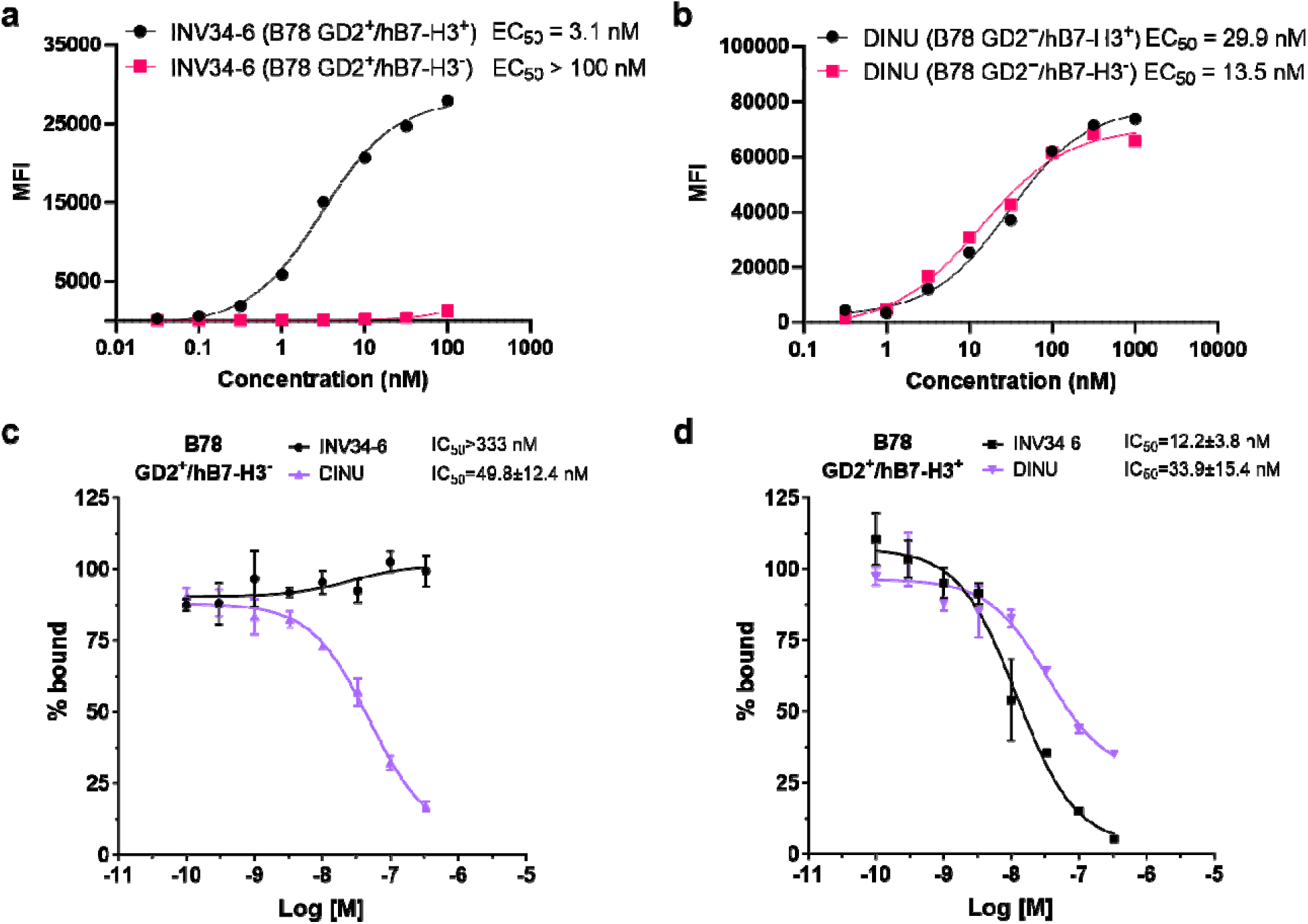
In vitro binding characterization of the avidity and selectivity for the bispecific GD2-B7-H3 targeting antibody. (**a**) INV34-6 showed selective binding to GD2^+^/hB7-H3^+^ B78 cells with high avidity compared to GD2^+^/hB7-H3^-^ B78 cells by flow cytometry. (**b**) Characterization of the binding properties for DINU using the differentially expressing GD2/hB7-H3 B78 cells showed high affinity to both GD2^+^ cell lines. Competition binding radioassays using (**c**) GD2^+^/hB7-H3^-^ B78 cells and (**d**) GD2^+^/hB7-H3^+^ B78 cells further demonstrated high avidity and selectivity of the bispecific GD2 and B7-H3 targeting platform.

Similar binding selectivity and avidity was observed for another bispecific antibody (INV36-6) using the B78 cells expressing both antigens (EC_50_ = 1 nM) or only GD2 (EC_50_ > 100 nM) (**Fig. S1**). Additional evaluation of bispecific antibody binding properties was conducted using competitive radioactive binding assays (**Fig. 2C-D**). For GD2^+^/hB7-H3^-^ B78 cells, no displacement of the radioactive probe was observed for INV34-6 (IC_50_ > 333 nM). However, binding was observed when INV34-6 was incubated with GD2^+^/hB7-H3^+^ B78 cells (IC_50_ of 49.8 ± 12.4 nM). Other bispecific antibodies targeting GD2 and B7-H3 were shown to have comparable binding properties (**Fig. S2**). No selectivity was observed for DINU, which bound to GD2^+^/hB7-H3^+^ B78 cells and GD2^+^/hB7-H3^-^ B78 cells with comparable affinities (IC_50_ = 33.9 ± 15.4 nM and IC_50_ = 49.8 ± 12.4 nM, respectively).

### Evaluating *in vivo* targeting using PET

We then investigated the *in vivo* targeting of INV34-6 radiolabeled with ^89^Zr (t_1/2_ = 74.2 h) in mice bearing GD2^+^/hB7-H3^+^ B78 tumors using PET. A non-targeted bispecific control antibody (bsAb CTRL) evaluated non-specific uptake contributions of the bispecific platform. Antibodies were conjugated and efficiently radiolabeled with ^89^Zr, as shown by the radiochemical yield of 94.3 ± 1.6% (n=4; **Fig. S3**) for [^89^Zr]Zr-Df-INV34-6. After conjugation, radiolabeling, and purification, PET images were acquired ([^89^Zr]Zr-Df-INV34-6 or [^89^Zr]Zr-Df-bsAb CTRL) in GD2^+^/hB7-H3^+^ tumor bearing mice (**Fig. 3A**; [^89^Zr]Zr-Df-INV33-2 or [^89^Zr]Zr-Df-INV36-6 in **Fig. S4; Tables S1-S2**). As shown in the maximum intensity projection (MIP) images, tumors can clearly be delineated using [^89^Zr]Zr-Df-INV34-6 as soon as 24 h post-injection (p.i.), whereas minimal tumor uptake was noted for non-specific [^89^Zr]Zr-Df-bsAb CTRL. The differences in the tumor and normal tissue uptake for [^89^Zr]Zr-Df-INV34-6 and [^89^Zr]Zr-Df-bsAb CTRL were quantified using volume of interest (VOI) analysis (**Fig. 3B**).

**Figure 3.**
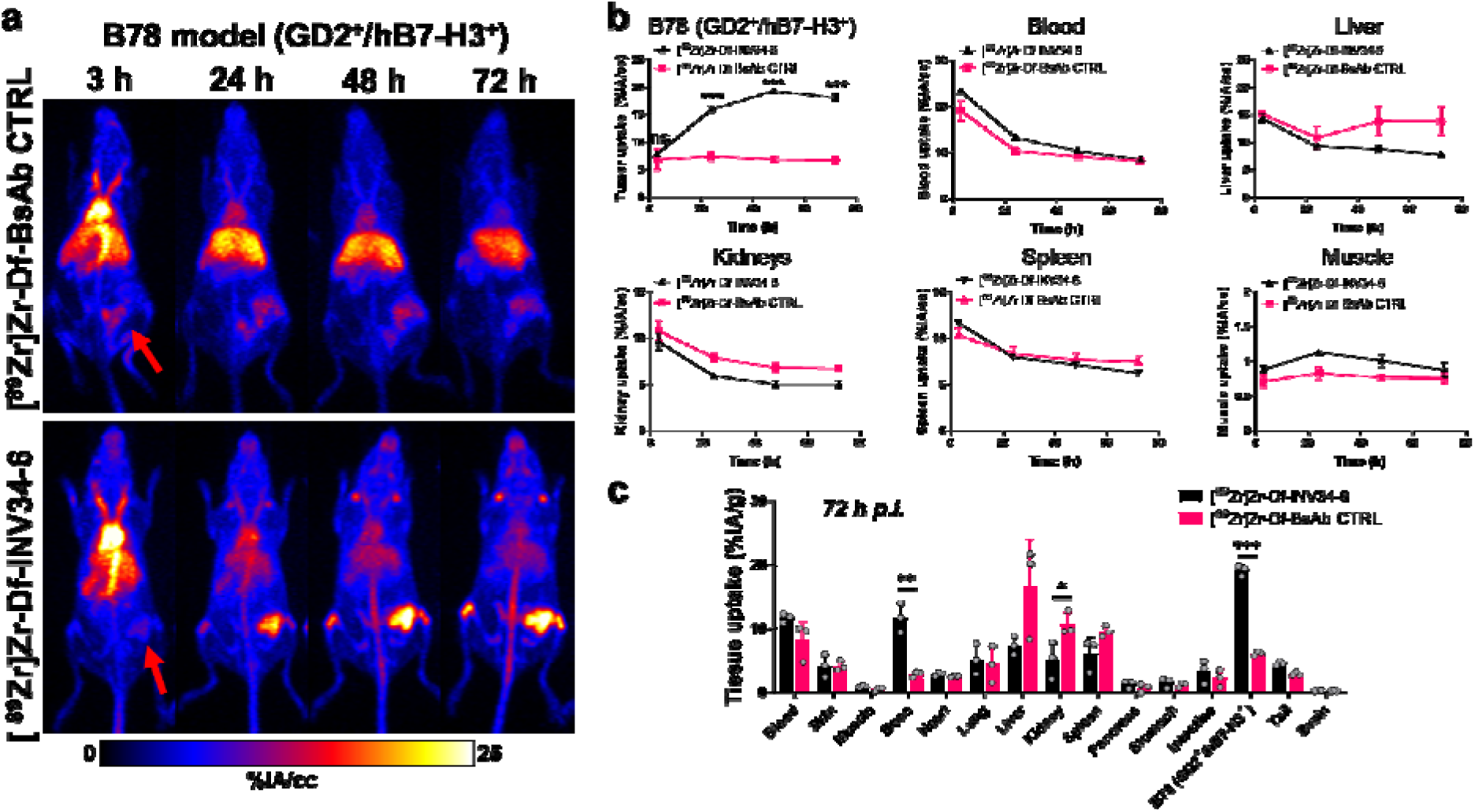
*In vivo* PET shows bispecific antibody targeting GD2-B7-H3 has high avidity for GD2^+^/B7H3^+^ B78 tumors compared to nonspecific control. (**a**) Representative maximum intensity projection PET images showed high tumor uptake for [^89^Zr]Zr-Df-INV34-6 compared to a non-specific control ([^89^Zr]Zr-Df-bsAb CTRL) in a GD2^+^/hB7-H3^+^ B78 tumor model. Red arrows show tumor location. (**b**) Volume of interest analysis found significantly higher tumor uptake and comparable normal tissue uptake for [^89^Zr]Zr-Df-INV34-6 following the initial imaging timepoint compared to [^89^Zr]Zr-Df-bsAb CTRL (n=3). (**c**) *Ex vivo* biodistribution studies confirmed higher uptake of [^89^Zr]Zr-Df-INV34-6 in the GD2^+^/hB7-H3^+^ B78 tumor model (n=3). p values (b-c) calculated using two-tailed Student’s t-test at each timepoint (ns = non-significant, *p < 0.05, **p < 0.01 ***p < 0.001).

Significantly higher tumor uptake for [^89^Zr]Zr-Df-INV34-6 was first observed at 24 h p.i. (16.0 ± 1.0 %IA/cc vs. 7.4 ± 1.4 %IA/cc; p = 0.00095) compared to [^89^Zr]Zr-Df-bsAb CTRL (7.4 ± 1.4 %IA/cc). The tumor uptake for [^89^Zr]Zr-Df-INV34-6 peaked at 48 h p.i. (19.3 ± 0.7 %IA/cc) and then plateaued at 72 h p.i. (18.2 ± 1 %IA/cc), which remained significantly higher than [^89^Zr]Zr-Df-bsAb CTRL at both timepoints (48 h: 6.9 ± 0.9 %IA/cc, p < 0.001; 72 h: 6.8 ± 1 %IA/cc, p < 0.001). The *in vivo* clearance of [^89^Zr]Zr-Df-INV34-6 and [^89^Zr]Zr-Df-bsAb CTRL from the blood was comparable over the PET study, with half-lives calculated using a one-phase exponential decay model to be 41.1 ± 2.7 h for [^89^Zr]Zr-Df-INV34-6 and 55.2 ± 27.4 h for [^89^Zr]Zr-Df-bsAb CTRL (p=0.42). In the other normal tissues of intestest (blood, liver, spleen, kidney, muscle), the uptake of [^89^Zr]Zr-Df-INV34-6 and [^89^Zr]Zr-Df-bsAb CTRL was comparable at all timepoints. Notably, higher uptake of [^89^Zr]Zr-Df-bsAb CTRL in the liver was observed albeit not significant. PET with other bispecific antibodies, [^89^Zr]Zr-Df-INV33-2 or [^89^Zr]Zr-Df-INV36-6, showed similar tumor and normal tissue uptake as [^89^Zr]Zr-Df-INV34-6 (**Fig. S5**), demonstrating the versatility of the bispecific platform.

Following the 72 h terminal imaging timepoint, the mice were euthanized, and the major tissues of interest were collected for *ex vivo* quantification (**Fig. 3C: Table S3**). The enhanced avidity of [^89^Zr]Zr-Df-INV34-6 was further demonstrated by significantly higher tumor uptake (19.1 ± 0.8 %IA/g) compared to [^89^Zr]Zr-Df-bsAb CTRL (6.1 ± 0.3 %IA/g, p < 0.001). Otherwise, the sole noteworthy difference in tissue uptake was observed in the bone, which was significantly higher (p = 0.003) for [^89^Zr]Zr-Df-INV34-6 (11.7 ± 2.4 %IA/g) compared to [^89^Zr]Zr-Df-bsAb CTRL (2.9 ± 0.4 %IA/g). No notable differences in the *ex vivo* quantification of the tumor uptake was found between the other GD2 and B7-H3 targeting bispecific antibodies of interest (**Fig. S6**). Overall, the biodistribution study corroborated the VOI analysis and demonstrated remarkable *in vivo* avidity toward GD2^+^/hB7-H3^+^ tumors.

### Evaluating *in vivo* selectivity using PET/CT

The selectivity of INV34-6 towards GD2+/hB7-H3^+^ cells was then compared to DINU. As such, we used PET to evaluate uptake of [^89^Zr]Zr-Df-INV34-6 and [^89^Zr]Zr-Df-DINU in either GD2^+^/hB7-H3^+^ or GD2^+^/hB7-H3 B78 tumor models. Representative MIP PET images of [^89^Zr]Zr-Df-INV34-6 and [^89^Zr]Zr-Df-DINU in GD2^+^/hB7-H3^+^ B78 tumors and GD2^+^/hB7-H3^-^ B78 tumors are shown in **Fig. 4**. VOI quantification of [^89^Zr]Zr-Df-INV34-6 and [^89^Zr]Zr-Df-DINU in mice bearing GD2^+^/hB7-H3^+^ or GD2^+^/hB7-H3^-^ B78 tumors showed similar biodistribution in normal tissues (**Fig. 5a-f; Table S4-S9**). Importantly, VOI quantification (**Fig. 5g; Table S10**) of [^89^Zr]Zr-Df-INV34-6 showed selectivity toward GD2^+^/hB7-H3^+^ B78 tumors compared to GD2^+^/hB7-H3^-^ B78 tumors, with increased relative tumor uptake at 48 h p.i. and 72 h p.i. (10.8 ± 2.4 %IA/cc vs. 7.3 ± 1.5 %IA/cc and 12.4 ± 1.9 %IA/cc vs. 7.7 ± 1.4 %IA/cc, respectively). The peak tumor uptake of [^89^Zr]Zr-Df-INV34-6 in GD2^+^/hB7-H3^+^ B78 tumors occurred at 72 h p.i.. In comparison, no significant difference in tumor uptake of [^89^Zr]Zr-Df-DINU was observed in GD2^+^/hB7-H3^+^ vs GD2^+^/hB7-H3^-^ B78 tumors. Furthermore, no significant differences in tumor uptake for [^89^Zr]Zr-Df-INV34-6 and [^89^Zr]Zr-Df-DINU in GD2^+^/hB7-H3^+^ B78 tumors were found at any timepoint. *Ex vivo* tissue quantification at 72 h p.i. further confirmed increased (p = 0.003) uptake of [^89^Zr]Zr-Df-INV34-6 in GD2^+^/hB7-H3^+^ B78 tumors (23.0 ± 2.4 %IA/g) compared to GD2^+^/hB7-H3^-^ B78 tumors (12.9 ± 1.4 %IA/g) (**Fig. 5h; Table S11**), while no significant difference was noted between the uptake of these antibodies in normal tissues (**Fig. S7**). No difference in [^89^Zr]Zr-Df-DINU tumor uptake was observed in GD2^+^/hB7-H3^-^ B78 (20.6 ± 3.4 %IA/g) and GD2^+^/hB7-H3^+^ B78 (21.1 ± 2.3 %IA/g) models.

**Figure 4.**
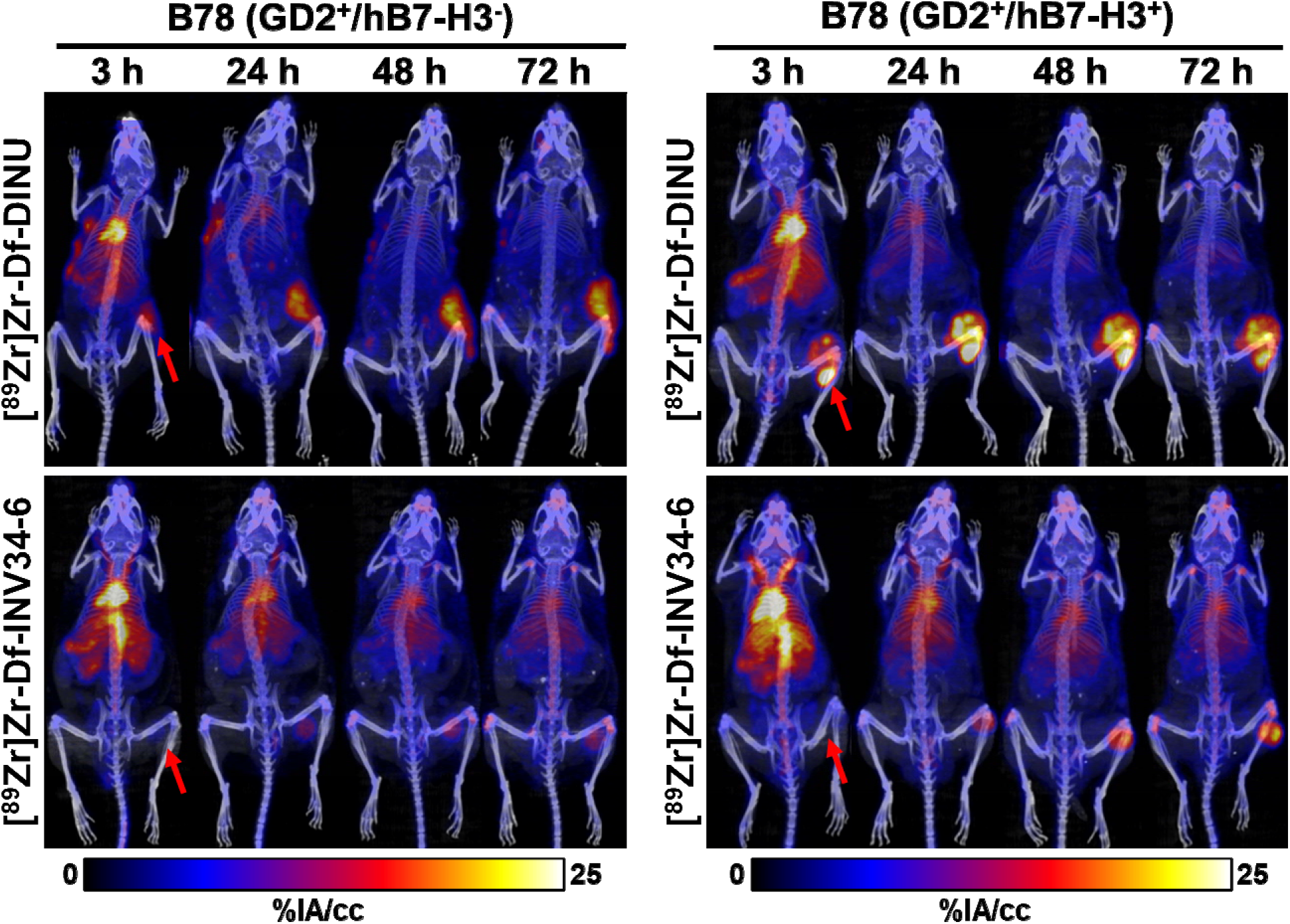
PET shows selectivity of GD2-B7H3 bispecific antibody *in vivo* compared to dinutuximab. Representative MIP PET images using radiolabeling [^89^Zr]Zr-Df-INV34-6 showed increased uptake in B78 GD2^+^/hB7-H3^+^ tumors compared to GD2^+^/B7-H3^-^ B78 tumors. In comparison, tumor uptake of [^89^Zr]Zr-Df-DINU in GD2^+^/hB7-H3^-^ B78 tumors and GD2^+^/hB7-H3^+^ B78 tumors was comparable. Red arrows indicate tumor location.

**Figure 5.**
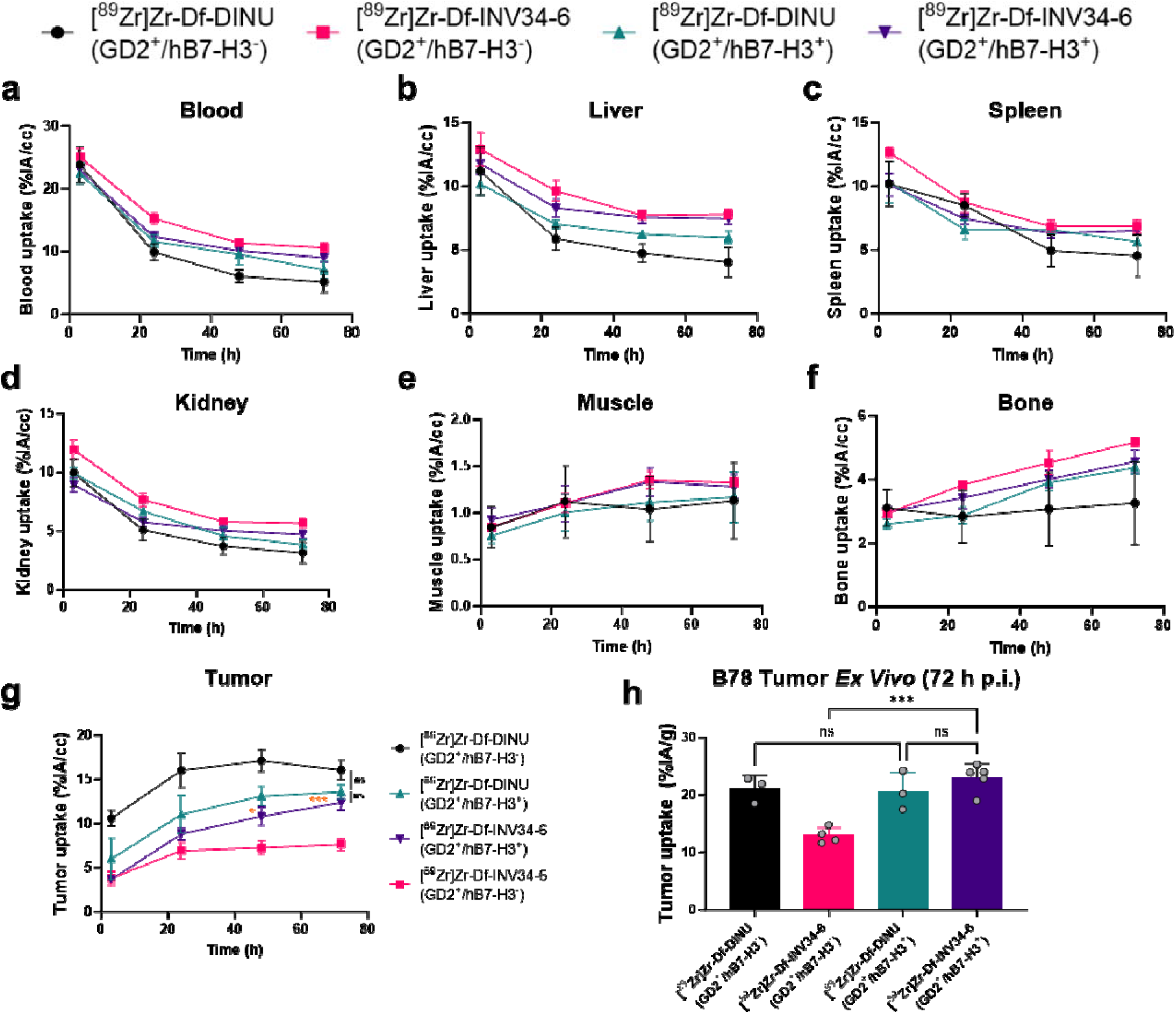
PET quantification of B78 tumor model with differential B7-H3 expression. Volume of interest quantification of [^89^Zr]Zr-Df-INV34-6 and [^89^Zr]Zr-Df-DINU in the (**a**) blood, (**b**) liver, (**c**) spleen, (**d**) kidney, (**e**) muscle, (**f**) bone, and (**g**) tumor at the imaging timepoints (%IA/cc) [n = 3-5]. (**h**) *Ex vivo* tumor uptake of [^89^Zr]Zr-Df-INV34-6 and [^89^Zr]Zr-Df-DINU in GD2^+^/hB7-H3^+^ and GD2^+^/hB7-H3^-^ B78 tumors (n=3-5). Orange p values compare [^89^Zr]Zr-Df-INV34-6 in GD2^+^/hB7-H3^+^ and GD2^+^/hB7-H3^-^ B78 tumors. p values were calculated using (g) two-tailed Student’s t-test at each timepoint or (h) one-way ANOVA with Tukey’s honest significant different post-hoc test (ns = non-significant at all timepoints, *p < 0.05. ***p < 0.001).

Moreover, no uptake difference was found between [^89^Zr]Zr-Df-DINU and [^89^Zr]Zr-Df-INV34-6 in GD2^+^/hB7-H3^+^ B78 tumors.

## Discussion

Bispecific antibodies have been proposed as a next generation therapeutic approach for targeted cancer treatment. While monoclonal antibodies are typically designed to target a single epitope on an antigen of interest, bispecific antibodies are engineered to target two separate antigen or epitopes. This dual-targeting approach enables more precise and potent antibody therapies against cancer cells while sparing undesired toxicities and potentially overcoming resistance mechanisms. Although bispecific antibodies are commonly investigated as cancer treatment, they are also assessed as treatments in autoimmune diseases and infectious diseases, among others.[17] The evolving landscape of bispecific antibodies in biopharmaceuticals underscores their importance as innovative tools that can revolutionize the therapeutic arsenal, providing novel avenues for personalized medicine and improving patient outcomes.

Antibody therapeutics targeting GD2 have been investigated in preclinically and clinically, becoming part of the standard of care for children with high-risk neuroblastoma[5, 18]. The expression of GD2 on the surface of cancer cells is associated with neuroblastoma, melanoma, osteosarcoma, small cell lung cancer, among others.[19] When anti-GD2 directed antibodies were initially utilized in the treatment of melanoma or neuroblastoma significant dose limiting toxicities stemming from GD2 expression on peripheral nerves was noted.[20, 21] Despite the undesirable side effects, anti-GD2 therapeutic antibodies improved overall and progression free survival rates when given as a treatment with granulocyte macrophage-colony stimulating factor and interleukin-2.[5, 18] Clinical studies using revised anti-GD2 antibody treatments, such as humanization, chimerization, combination therapies, and Fc mutations, have continued to observe acute, severe pain from treatment.[22–25] We propose anti-GD2 directed treatment would benefit from a requirement of co-expression of another marker, B7-H3, using the bispecific platform with turned affinities. Importantly, B7-H3 expression is not found on nerves, which may minimize the pain toxicity associated with GD2-targeting therapies.[26] B7-H3 is considered an immune checkpoint molecule and pan-cancer antigen with a role in disease progression and metastasis.[27–29] Studies have found high B7-H3 expression in both melanoma and neuroblastoma patient populations that correlated with poor survival outcomes.[30–32] Considering the constraints associated with anti-GD2 targeting biologics, we suggest a novel approach centered around conditional B7-H3 expression. This could potentially revolutionize the therapeutic options available for patients targeted by these treatments.

Our study explored the potential of a bispecific antibody platform to increase the selectivity toward targeting cells expressing GD2 and B7-H3. Our innovative approach was validated through a series of cell binding and PET studies utilizing B78 melanoma cells with differential GD2/B7-H3 expression profiles. The results of these studies demonstrated that the GD2 and B7-H3 targeting bispecific antibodies selectively bind to cancer cells expressing both GD2 and B7-H3 with high avidity, showcasing their potential to precisely target tumor cells while sparing normal cells (i.e., peripheral nerves) expressing only GD2.

A similar dual-targeting approach using GD2 and B7-H3 was demonstrated using chimeric antigen targeting (CAR) T-cell therapy.[33] This platform may avert the limitations of other CAR-T cell therapies, such as limited accessibility for patients, complex manufacturing processes, and poor efficacy in solid tumors, and also requires the use of non-human transcription factors.[34] In comparison, we have found that the bispecific platform can be manufactured at scale and easily purified using traditional antibody purification techniques. The improved selectivity demonstrated in this proof-of-concept study enables further investigations into potential therapies using this platform. Subsequent investigations may explore therapeutic avenues involving antibody-dependent cellular cytotoxicity, antibody-drug conjugates, or radioimmunotherapy. Using one or more of these therapeutic mechanisms of actions, a bispecific antibody requiring simultaneous expression of GD2 and B7-H3 may improve treatment outcomes and quality of life for patients of many cancer types.

The advancements we demonstrated for targeting GD2-positive tumors using a GD2 and B7-H3 targeting bispecific antibody has limitations. Our proof-of-concept study relied on engineering a murine melanoma cell line to establish differential B7-H3 and GD2 status. However, it should be acknowledged that species differences, such as receptor densities, necessitate evaluation using clinically relevant human cell lines or patient derived xenografts models. Additionally, we limited our study to using PET to demonstrate the enhanced selectivity toward cancers cells with our bispecific antibody strategy. It is imperative to assess whether the improved selectivity translates into improved treatment efficacy in subsequent studies. Furthermore, even though these studies here show minimal binding of bispecific antibodies to cells expressing GD2 in the absence of B7-H3, further studies evaluating actual binding of these bispecific antibodies (in comparison to dinutuximab) to nerves, and their relative induction of pain in vivo are still required. Additional validation of our findings that bispecific antibodies improve antibody selectivity should be extended to other tumors expressing GD2 and B7-H3. Analyses addressing these issues are beyond the scope of this manuscript, but several of these questions are addressed in a separate manuscript evaluating a novel anti-GD2/anti-B7-H3 bispecific antibody [35]. Following the necessary preclinical studies, the next generation GD2/B7-H3 bispecific antibody holds promise to supersede existing anti-GD2 antibody therapies for cancer patients, ultimately improving treatment quality.

## Conclusion

We designed a bispecific antibody to target GD2 and B7-H3 in a manner that requires co-expression on a single cell. By doing so, the selectivity of antibody binding was engineered toward GD2^+^/hB7-H3^+^ cells over GD2^+^/hB7-H3^-^ and resulting in high avidity, improving cancer cell targeting and potentially limiting side effects attributed to binding non targeted tissues such as nerves expressing GD2 only. Binding assays showed that INV34-6 displayed high avidity and selectivity to GD2^+^/hB7-H3^+^ over GD2^+^/hB7-H3^-^ B78 cells, whereas DINU bound similarly to both GD2 expressing cell lines. PET showed that the targeting improvements for INV34-6 over DINU or a non-specific control (bsAb CTRL) were retained *in vivo* using GD2^+^/hB7-H3^+^ and GD2^+^/hB7-H3^-^ B78 tumor models. Our results demonstrate that tuning the avidity toward antigens using bispecific antibodies is an effective strategy to improve selective targeting of cancer and may improve cancer therapies. PET was fundamental in showing that our antibody engineering approach translated into improved selectivity *in vivo*.

## Question

Can a GD2 and B7-H3 targeting bispecific antibody be used to enhance the selectivity toward GD2 positive tumors?

## Pertinent Findings

The bispecific antibody showed high avidity and selectivity toward tumors expressing GD2 and B7-H3.

## Implications for patient care

Requiring dual expression of GD2 and B7-H3 using a bispecific antibody may provide a next-generation therapeutic approach to limit neuropathic pain caused by existing GD2 targeting antibodies.

## Acknowledgments

This work was supported by the University of Wisconsin-Madison, the National Institutes of Health (R01HL153721, P01CA250972, R35CA197078, T32CA009206) and Department of Defense (Early Investigator Award, W81XWH1910285), The Wisconsin Alumni Research Foundation, Midwest Athletes Against Childhood Cancer; the University of Wisconsin Carbone Cancer Center and research grants from the Pablove Foundation, the HESI-thrive Foundation, the Hyundai Hope on Wheels Foundation, and the End Kids Cancer Foundation. The content is solely the responsibility of the authors and does not necessarily represent the official views of the National Institutes of Health. The authors wish to acknowledge the Small Animal Imaging and Radiotherapy Facility (SAIRF) at UW-Madison maintaining facilities for acquiring PET/CT, including support through the Cancer Center Support Grant NCI P30CA014520), and Invenra Inc. (Madison WI) for providing the bispecific mAbs used in this research.

