## Supplemental Information for "Targeting both GD2 and B7-H3 using bispecific antibody improves tumor selectivity for GD2-positive tumors"

**Running title:** Bispecific antibodies improve selectivity for GD2-positive tumors

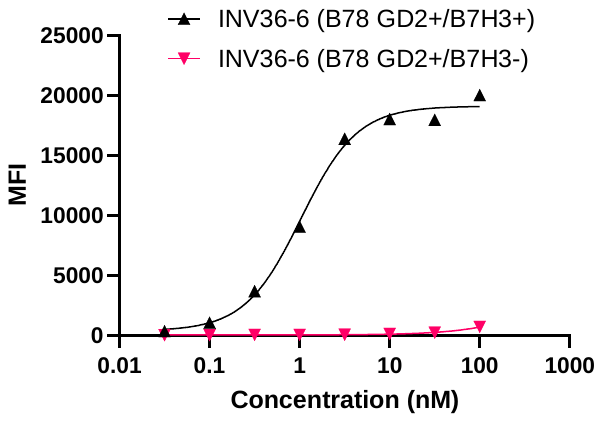

**Figure S1.** Cell binding assay for other bispecific body clone (INV36-6) targeting GD2 and B7-H3.

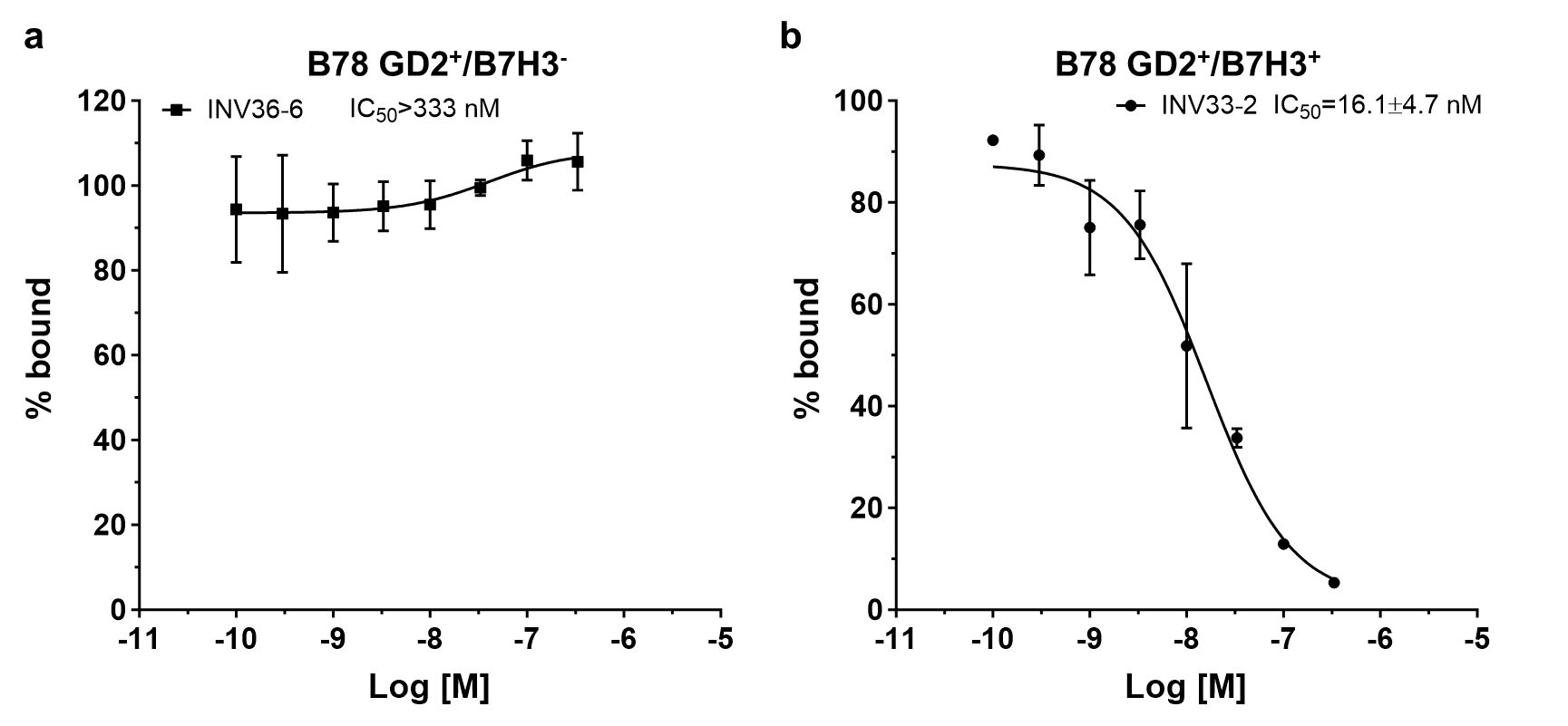

**Figure S2.** Competition binding assays using (A) B78 GD2+/B7-H3- cells or (B) B78 GD2+/B7-H3+ cells show that other bispecific antibodies targeting GD2 and B7-H3 maintain selectivity and avidity for cancer cells expressing both targets.

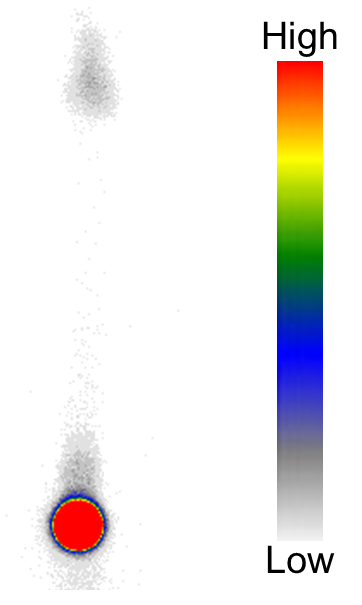

**Figure S3.** Representative radio-TLC showing highly efficiency Zr-89 radiolabeling of deferoxamine (Df) conjugated antibodies.

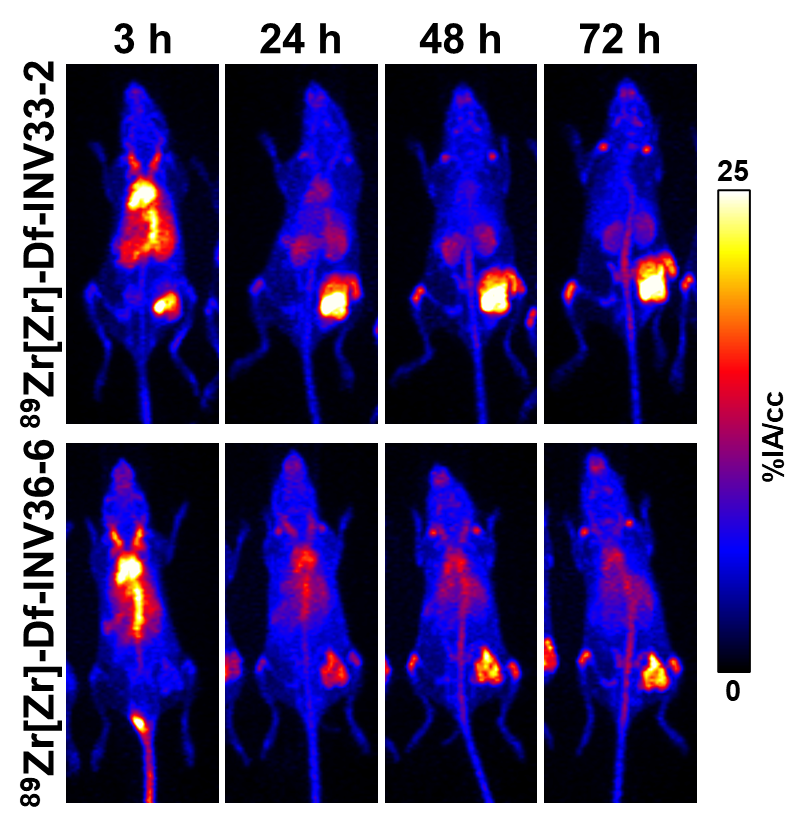

**Figure S4.** Representative maximum intensity projection PET images of ^89^Zr[Zr]-Df-INV33-2 and ^89^Zr[Zr]-Df-INV36-6 targeting GD2 and B7-H3 in the B78 GD2^+^/B7-H3^+^ tumor model.

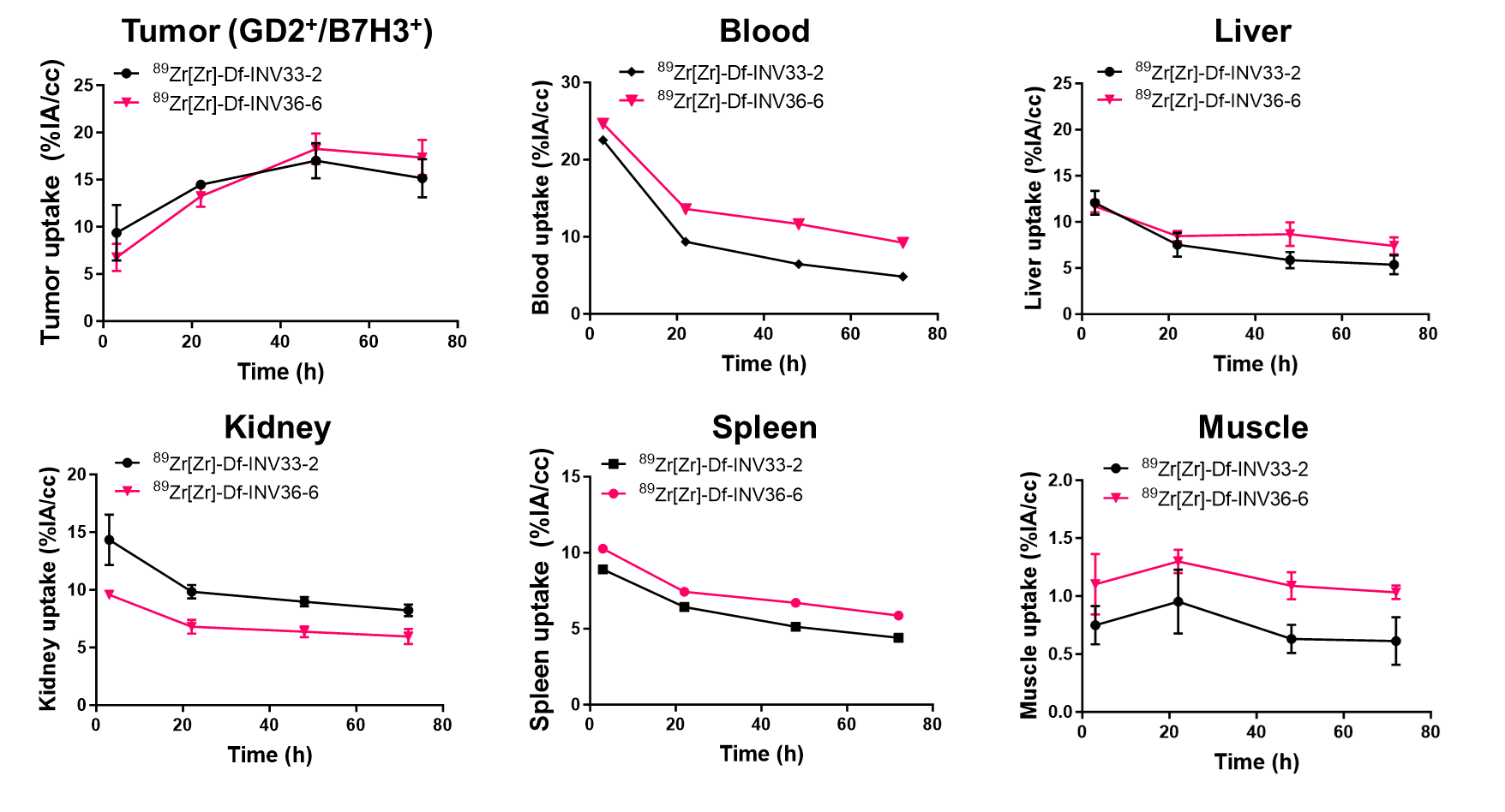

**Figure S5.** Volume of interest quantification for ^89^Zr[Zr]-Df-INV33-2 and ^89^Zr[Zr]-Df-INV36-6 in the B78 GD2^+^/B7-H3^+^ tumor model (n=3).

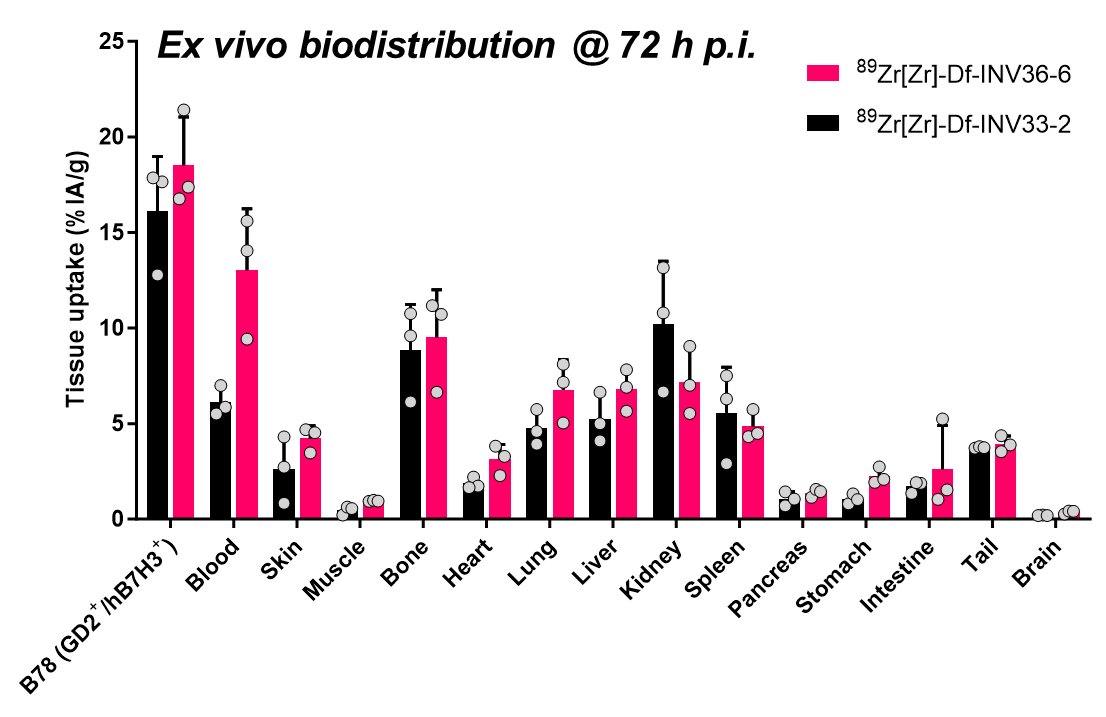

**Figure S6.** Ex vivo quantification for ^89^Zr[Zr]-Df-INV33-2 and ^89^Zr[Zr]-Df-INV36-6 in the B78 GD2^+^/B7-H3^+^ tumor model (n=3).

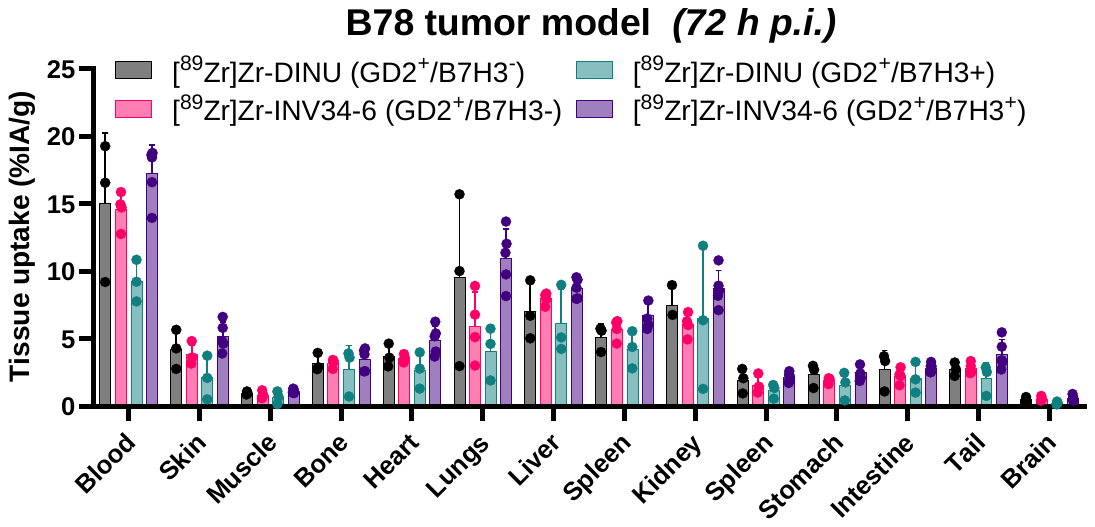

**Figure S7.** Ex vivo quantification for ^89^Zr[Zr]-Df-INV34-6 and ^89^Zr[Zr]-Df-DINU in the GD2^+^/B7-H3^+^ B78 tumor model and GD2^+^/B7-H3^-^ B78 tumor model (n=3-5).

**Table S1.** Volume of Interest (VOI) Quantification for [^89^Zr]Zr-Df-bsAb CTRL in the GD2^+^/hB7-H3^+^ B78 tumor model (n=3).

| [^89^Zr]Zr-Df-bsAb CTRL | | | | | | | | | | | | |
| --- | --- | --- | --- | --- | --- | --- | --- | --- | --- | --- | --- | --- |
| Time (h) | Heart (%IA/cc) | | Liver (%IA/cc) | | Kidneys (%IA/cc) | | Spleen (%IA/cc) | | Muscle (%IA/cc) | | Tumor (%IA/cc) | |
|  | AVG | SD | AVG | SD | AVG | SD | AVG | SD | AVG | SD | AVG | SD |
| 3 | 19.2 | 3.8 | 15.3 | 0.5 | 10.8 | 1.8 | 10.4 | 1.3 | 0.7 | 0.1 | 6.9 | 3.4 |
| 24 | 10.5 | 0.6 | 10.9 | 3.5 | 7.9 | 0.7 | 8.4 | 1.2 | 0.8 | 0.2 | 7.4 | 1.4 |
| 48 | 9.2 | 0.7 | 14.0 | 4.5 | 6.8 | 0.9 | 7.7 | 1.1 | 0.8 | 0.0 | 6.9 | 0.9 |
| 72 | 8.3 | 0.5 | 13.9 | 4.4 | 6.8 | 0.3 | 7.5 | 1.1 | 0.8 | 0.1 | 6.8 | 1.0 |

**Table S2.** Volume of interest (VOI) quantification for [^89^Zr]Zr-Df-INV34-6 in the GD2^+^/hB7-H3^+^ B78 tumor model (n=3).

| [^89^Zr]Zr-Df-INV34-6 | | | | | | | | | | | | |
| --- | --- | --- | --- | --- | --- | --- | --- | --- | --- | --- | --- | --- |
| Time (h) | Heart (%IA/cc) | | Liver (%IA/cc) | | Kidneys (%IA/cc) | | Spleen (%IA/cc) | | Muscle (%IA/cc) | | Tumor (%IA/cc) | |
|  | AVG | SD | AVG | SD | AVG | SD | AVG | SD | AVG | SD | AVG | SD |
| 3 | 23.4 | 1.0 | 14.4 | 1.1 | 9.6 | 1.6 | 11.7 | 0.9 | 0.9 | 0.1 | 7.9 | 0.7 |
| 24 | 13.3 | 0.8 | 9.4 | 0.8 | 6.0 | 0.3 | 8.0 | 0.6 | 1.1 | 0.1 | 16.0 | 1.0 |
| 48 | 10.4 | 0.6 | 8.8 | 1.0 | 5.0 | 0.6 | 7.2 | 1.0 | 1.0 | 0.2 | 19.3 | 0.7 |
| 72 | 8.6 | 0.8 | 7.8 | 0.6 | 5.0 | 0.7 | 6.3 | 0.6 | 0.9 | 0.2 | 18.2 | 1.0 |

**Table S3.** Ex vivo quantification for the biodistribution of [^89^Zr]Zr-Df-bsAb CTRL and [^89^Zr]Zr-Df-INV34-6 in the GD2^+^/hB7-H3^+^ B78 tumor model (n=3).

| Tissue uptake (%IA/g) | [^89^Zr]Zr-Df- bsAb CTRL | | [^89^Zr]Zr-Df-INV34-6 | |
| --- | --- | --- | --- | --- |
|  | AVG | SD | AVG | SD |
| Blood | 8.1 | 3.0 | 11.7 | 0.7 |
| Skin | 4.1 | 0.8 | 4.2 | 1.8 |
| Muscle | 0.5 | 0.3 | 0.9 | 0.1 |
| Bone | 2.9 | 0.4 | 11.7 | 2.4 |
| Heart | 2.5 | 0.1 | 2.9 | 0.3 |
| Lung | 4.4 | 2.9 | 5.2 | 2.4 |
| Liver | 16.7 | 7.3 | 7.3 | 1.5 |
| Kidney | 10.7 | 1.9 | 5.1 | 2.9 |
| Spleen | 9.6 | 0.9 | 6.1 | 2.6 |
| Pancreas | 0.7 | 0.6 | 1.3 | 0.6 |
| Stomach | 1.1 | 0.4 | 1.5 | 0.8 |
| Intestine | 2.2 | 1.4 | 3.3 | 1.8 |
| B78 tumor (GD2^+^/hB7-H3^+^) | 6.1 | 0.3 | 19.1 | 0.8 |
| Tail | 2.8 | 0.3 | 4.5 | 0.3 |
| Brain | 0.27 | 0.04 | 0.3 | 0.1 |

**Table S4.** Volume of interest (VOI) quantification for [^89^Zr]Zr-Df-DINU and [^89^Zr]Zr-Df-DINU in the blood of the GD2^+^/hB7-H3^-^  and GD2^+^/hB7-H3^+^ B78 tumor models (n=3-5).

| Blood uptake (%IA/cc) | [^89^Zr]Zr-Df-DINU (GD2^+^/hB7-H3^-^) | | [^89^Zr]Zr-Df-INV34-6 (GD2^+^/hB7-H3^-^) | | | [^89^Zr]Zr-Df-DINU (GD2^+^/hB7-H3^+^) | | | [^89^Zr]Zr-Df-INV34-6 (GD2^+^/hB7-H3^+^) | |
| --- | --- | --- | --- | --- | --- | --- | --- | --- | --- | --- |
| Time (h) | AVG | SD | AVG | SD | AVG | | SD | AVG | | SD |
| 3 | 23.8 | 5.0 | 25.1 | 2.7 | 22.5 | | 2.9 | 23.0 | | 1.9 |
| 24 | 9.9 | 2.3 | 15.2 | 1.7 | 11.6 | | 1.8 | 12.3 | | 1.8 |
| 48 | 6.1 | 1.8 | 11.4 | 1.2 | 9.5 | | 2.9 | 10.1 | | 2.0 |
| 72 | 5.1 | 3.0 | 10.6 | 1.6 | 7.0 | | 2.4 | 9.0 | | 1.6 |

**Table S5.** Volume of interest (VOI) quantification for [^89^Zr]Zr-Df-DINU and [^89^Zr]Zr-Df-DINU in the liver of the GD2^+^/hB7-H3^-^  and GD2^+^/hB7-H3^+^ B78 tumor models (n=3-5).

| Liver uptake (%IA/cc) | [^89^Zr]Zr-Df-DINU (GD2^+^/hB7-H3^-^) | | [^89^Zr]Zr-Df-INV34-6 (GD2^+^/hB7-H3^-^) | | | [^89^Zr]Zr-Df-DINU (GD2^+^/hB7-H3^+^) | | | [^89^Zr]Zr-Df-INV34-6 (GD2^+^/hB7-H3^+^) | |
| --- | --- | --- | --- | --- | --- | --- | --- | --- | --- | --- |
| Time (h) | AVG | SD | AVG | SD | AVG | | SD | AVG | | SD |
| 3 | 11.2 | 3.2 | 12.9 | 2.6 | 10.2 | | 1.6 | 11.7 | | 1.9 |
| 24 | 5.9 | 1.5 | 9.6 | 1.7 | 7.0 | | 0.6 | 8.3 | | 1.5 |
| 48 | 4.7 | 1.3 | 7.7 | 0.7 | 6.2 | | 0.5 | 7.6 | | 1.0 |
| 72 | 4.0 | 2.0 | 7.8 | 0.8 | 6.0 | | 0.9 | 7.4 | | 1.0 |

**Table S6.** Volume of interest (VOI) quantification for [^89^Zr]Zr-Df-DINU and [^89^Zr]Zr-Df-DINU in the spleen of the GD2^+^/hB7-H3^-^  and GD2^+^/hB7-H3^+^ B78 tumor models (n=3-5).

| Spleen uptake (%IA/cc) | [^89^Zr]Zr-Df-DINU (GD2^+^/hB7-H3^-^) | | [^89^Zr]Zr-Df-INV34-6 (GD2^+^/hB7-H3^-^) | | | [^89^Zr]Zr-Df-DINU (GD2^+^/hB7-H3^+^) | | | [^89^Zr]Zr-Df-INV34-6 (GD2^+^/hB7-H3^+^) | |
| --- | --- | --- | --- | --- | --- | --- | --- | --- | --- | --- |
| Time (h) | AVG | SD | AVG | SD | AVG | | SD | AVG | | SD |
| 3 | 10.2 | 3.0 | 12.7 | 0.8 | 10.3 | | 2.8 | 10.1 | | 2.0 |
| 24 | 8.5 | 1.6 | 8.7 | 1.7 | 6.6 | | 1.3 | 7.4 | | 0.9 |
| 48 | 4.9 | 2.1 | 6.8 | 1.0 | 6.6 | | 0.4 | 6.3 | | 0.9 |
| 72 | 4.5 | 2.8 | 6.9 | 1.0 | 5.6 | | 0.5 | 6.5 | | 0.8 |

**Table S7.** Volume of interest (VOI) quantification for [^89^Zr]Zr-Df-DINU and [^89^Zr]Zr-Df-DINU in the kidney of the GD2^+^/hB7-H3^-^  and GD2^+^/hB7-H3^+^ B78 tumor models (n=3-5).

| Kidney uptake (%IA/cc) | [^89^Zr]Zr-Df-DINU (GD2^+^/hB7-H3^-^) | | [^89^Zr]Zr-Df-INV34-6 (GD2^+^/hB7-H3^-^) | | | [^89^Zr]Zr-Df-DINU (GD2^+^/hB7-H3^+^) | | | [^89^Zr]Zr-Df-INV34-6 (GD2^+^/hB7-H3^+^) | |
| --- | --- | --- | --- | --- | --- | --- | --- | --- | --- | --- |
| Time (h) | AVG | SD | AVG | SD | AVG | | SD | AVG | | SD |
| 3 | 10.0 | 2.0 | 12.0 | 1.6 | 9.9 | | 0.9 | 9.0 | | 1.3 |
| 24 | 5.1 | 1.6 | 7.7 | 1.2 | 6.7 | | 1.3 | 5.8 | | 0.9 |
| 48 | 3.7 | 1.3 | 5.8 | 0.5 | 4.6 | | 1.2 | 5.1 | | 1.0 |
| 72 | 3.2 | 1.6 | 5.7 | 0.7 | 3.9 | | 0.8 | 4.7 | | 0.8 |

**Table S8.** Volume of interest (VOI) quantification for [^89^Zr]Zr-Df-DINU and [^89^Zr]Zr-Df-DINU in the muscle of the GD2^+^/hB7-H3^-^  and GD2^+^/hB7-H3^+^ B78 tumor models (n=3-5).

| Muscle uptake (%IA/cc) | [^89^Zr]Zr-Df-DINU (GD2^+^/hB7-H3^-^) | | [^89^Zr]Zr-Df-INV34-6 (GD2^+^/hB7-H3^-^) | | | [^89^Zr]Zr-Df-DINU (GD2^+^/hB7-H3^+^) | | | [^89^Zr]Zr-Df-INV34-6 (GD2^+^/hB7-H3^+^) | |
| --- | --- | --- | --- | --- | --- | --- | --- | --- | --- | --- |
| Time (h) | AVG | SD | AVG | SD | AVG | | SD | AVG | | SD |
| 3 | 0.8 | 0.4 | 0.8 | 0.1 | 0.8 | | 0.1 | 0.9 | | 0.3 |
| 24 | 1.1 | 0.7 | 1.1 | 0.2 | 1.0 | | 0.3 | 1.1 | | 0.4 |
| 48 | 1.0 | 0.6 | 1.4 | 0.2 | 1.1 | | 0.3 | 1.3 | | 0.3 |
| 72 | 1.1 | 0.7 | 1.3 | 0.1 | 1.2 | | 0.5 | 1.3 | | 0.3 |

**Table S9.** Volume of interest (VOI) quantification for [^89^Zr]Zr-Df-DINU and [^89^Zr]Zr-Df-DINU in the bone of the GD2^+^/hB7-H3^-^  and GD2^+^/hB7-H3^+^ B78 tumor models (n=3-5).

| Bone uptake (%IA/cc) | [^89^Zr]Zr-Df-DINU (GD2^+^/hB7-H3^-^) | | [^89^Zr]Zr-Df-INV34-6 (GD2^+^/hB7-H3^-^) | | | [^89^Zr]Zr-Df-DINU (GD2^+^/hB7-H3^+^) | | | [^89^Zr]Zr-Df-INV34-6 (GD2^+^/hB7-H3^+^) | |
| --- | --- | --- | --- | --- | --- | --- | --- | --- | --- | --- |
| Time (h) | AVG | SD | AVG | SD | AVG | | SD | AVG | | SD |
| 3 | 3.1 | 1.0 | 2.9 | 0.3 | 2.6 | | 0.3 | 3.0 | | 0.4 |
| 24 | 2.8 | 1.4 | 3.8 | 0.3 | 2.9 | | 0.5 | 3.4 | | 0.8 |
| 48 | 3.1 | 2.0 | 4.5 | 0.8 | 3.9 | | 0.4 | 4.0 | | 0.6 |
| 72 | 3.3 | 2.2 | 5.2 | 0.3 | 4.4 | | 0.3 | 4.6 | | 0.8 |

**Table S10.** Volume of interest (VOI) quantification for [^89^Zr]Zr-Df-DINU and [^89^Zr]Zr-Df-DINU in the tumor of the GD2^+^/hB7-H3^-^  and GD2^+^/hB7-H3^+^ B78 tumor models (n=3-5).

| Tumor uptake (%IA/cc) | [^89^Zr]Zr-Df-DINU (GD2^+^/hB7-H3^-^) | | [^89^Zr]Zr-Df-INV34-6 (GD2^+^/hB7-H3^-^) | | | [^89^Zr]Zr-Df-DINU (GD2^+^/hB7-H3^+^) | | | [^89^Zr]Zr-Df-INV34-6 (GD2^+^/hB7-H3^+^) | |
| --- | --- | --- | --- | --- | --- | --- | --- | --- | --- | --- |
| Time (h) | AVG | SD | AVG | SD | AVG | | SD | AVG | | SD |
| 3 | 10.6 | 1.5 | 3.8 | 1.5 | 6.0 | | 4.1 | 3.7 | | 0.8 |
| 24 | 16.0 | 3.4 | 7.0 | 1.8 | 11.0 | | 3.7 | 8.8 | | 1.5 |
| 48 | 17.1 | 2.1 | 7.3 | 1.5 | 13.1 | | 1.9 | 10.8 | | 2.4 |
| 72 | 16.1 | 2.0 | 7.7 | 1.4 | 13.6 | | 1.5 | 12.4 | | 1.9 |

**Table S11.** Ex vivo quantification for the biodistribution of [^89^Zr]Zr-Df-DINU and [^89^Zr]Zr-Df-DINU in the GD2^+^/hB7-H3^-^  and GD2^+^/hB7-H3^+^ B78 tumor models (n=3-5).

| Tissue uptake (%IA/g) | [^89^Zr]Zr-DINU (GD2^+^/B7-H3^-^) | | [^89^Zr]Zr-INV34-6 (GD2^+^/B7-H3^-^) | | [^89^Zr]Zr-DINU (GD2^+^/B7-H3^+^) | | [^89^Zr]Zr-INV34-6 (GD2^+^/B7-H3^+^) | |
| --- | --- | --- | --- | --- | --- | --- | --- | --- |
|  | AVG | SD | AVG | SD | AVG | SD | AVG | SD |
| Blood | 15.02 | 5.21 | 14.60 | 1.31 | 9.30 | 1.55 | 17.29 | 2.05 |
| Skin | 4.25 | 1.45 | 3.84 | 0.71 | 2.13 | 1.62 | 5.18 | 1.06 |
| Muscle | 0.99 | 0.11 | 0.85 | 0.26 | 0.67 | 0.47 | 1.14 | 0.14 |
| Bone | 3.21 | 0.66 | 3.22 | 0.30 | 2.75 | 1.76 | 3.50 | 0.84 |
| Heart | 3.75 | 0.86 | 3.57 | 0.26 | 2.71 | 1.34 | 4.93 | 1.05 |
| Lungs | 9.58 | 6.38 | 5.98 | 2.50 | 4.10 | 1.97 | 11.02 | 2.12 |
| Liver | 7.03 | 2.16 | 7.98 | 0.44 | 6.13 | 2.53 | 8.75 | 0.74 |
| Spleen | 5.14 | 0.97 | 5.73 | 0.75 | 4.26 | 1.38 | 6.80 | 0.99 |
| Kidney | 7.52 | 1.28 | 6.07 | 0.84 | 6.53 | 5.31 | 8.74 | 1.35 |
| Spleen | 1.95 | 0.91 | 1.60 | 0.60 | 1.17 | 0.54 | 2.18 | 0.34 |
| Stomach | 2.37 | 0.88 | 1.94 | 0.19 | 1.58 | 1.04 | 2.52 | 0.54 |
| Intestine | 2.73 | 1.41 | 2.28 | 0.55 | 2.12 | 1.15 | 2.86 | 0.33 |
| Tail | 2.76 | 0.50 | 2.85 | 0.42 | 2.07 | 1.14 | 3.88 | 1.08 |
| Brain | 0.53 | 0.15 | 0.57 | 0.15 | 0.26 | 0.11 | 0.61 | 0.20 |
